# Chlorpheniramine Maleate Displays Multiple Modes of Antiviral Action Against SARS-CoV-2: A Mechanistic Study

**DOI:** 10.1101/2023.08.28.554806

**Authors:** Yaseen A. M. M. Elshaier, Ahmed Mostafa, Fernando Valerio-Pascua, Mari L. Tesch, Joshua M. Costin, Franck F. Rahaghi

## Abstract

Chlorpheniramine Maleate (CPM) has been identified as a potential antiviral compound against Severe Acute Respiratory Syndrome Coronavirus 2 (SARS-CoV-2). In this study, we investigated the in vitro effects of CPM on key stages of the SARS-CoV-2 replication cycle, including viral adsorption, replication inhibition, and virucidal activity. Our findings demonstrate that CPM exhibits antiviral properties by interfering with viral adsorption, replication, and directly inactivating the virus. Molecular docking analysis revealed interactions between CPM and essential viral proteins, such as the main protease receptor, spike protein receptor, and RNA polymerase. CPM’s interactions were primarily hydrophobic in nature, with an additional hydrogen bond formation in the RNA polymerase active site. These results suggest that CPM has the potential to serve as a multitarget antiviral agent against SARS-CoV-2 and potentially other respiratory viruses. Further investigations are warranted to explore its clinical implications and assess its efficacy in vivo.

## Introduction

Since the emergence of severe acute respiratory syndrome coronavirus 2 (SARS-CoV-2) in Wuhan, China, a few drugs have been approved for the treatment and prophylaxis of coronavirus disease 2019 (COVID-19). Various attempts have been made to identify and repurpose currently approved drugs for the treatment of COVID-19, specifically those with antiviral potentials such as the antihistamine drugs Hydroxyzine, Azelastine, Carbinoxamine maleate, and Chlorpheniramine maleate (CPM) (A. Mostafa et al., 2020; Reznikov et al., 2021; Xu et al., 2018). CPM is a first-generation H1 antihistamine, that has been a longstanding treatment option for allergies, hay fever, common cold, cough, and nasal congestion. Its anti-inflammatory and immunomodulatory properties likely contribute to its efficacy (Lee et al., 2014; Rizvi et al., 2022). It is worth noting that several studies have demonstrated CPM’s antiviral activity against respiratory viruses such as influenza and SARS-CoV-2 (Black, 2022; A. Mostafa et al., 2020; Westover et al., 2020; Xu et al., 2018). Furthermore, early clinical studies have indicated CPM’s effectiveness in treating COVID-19 (Black, 2022; Morán Blanco et al., 2021; Sanchez-Gonzalez et al., 2022; Torres et al., 2021). Our research team has also reported on the safety and efficacy of intranasally administered CPM for the treatment of COVID-19 and allergic rhinitis (Sanchez-Gonzalez et al., 2021; Sanchez-Gonzalez et al., 2022).

Mechanistically, Hydroxyzine, and related antihistamines can inhibit SARS-CoV-2 entry via off-target inhibitory ACE2 activity by forming intermolecular interactions with the active site and virus replication by binding the sigma-1 receptor (Reznikov et al., 2021). While the mode of action of CPM against influenza is well-established, the mechanisms underlying its antiviral effect against SARS-CoV-2, including the corresponding cellular compartments and molecular targets, remain unknown. Accordingly, this study aimed to determine CPM’s mode of antiviral action and molecular targets against SARS-CoV-2.

## Methods

### Cells and virus strain

Vero-E6 cells were cultured in Dulbecco’s modified Eagle’s medium (DMEM) (Invitrogen, Germany) containing 1 % Penicillin/Streptomycin mixture and 10 % Fetal Bovine Serum (FBS) in a humidified incubator at 37 °C and 5 % CO_2_. The hCoV-19/Egypt/NRC-3/2020 “NRC-03-nhCoV” virus, belonging to the ancestor clade A2a (Kandeil et al., 2020) was propagated and titrated as described previously (Mostafa et al., 2020).

### MTT cytotoxicity assay

To assess the half maximal cytotoxic concentration (CC50), stock solutions of the test compounds were prepared in 10 % Dimethyl sulfoxide (DMSO) in double-distilled water (ddH2O) and diluted further to the working solutions with DMEM. The cytotoxic activity of the drug CPM was tested in cells by using the 3 4-(, 5-dimethylthiazol -2-yl)-2, 5-diphenyltetrazolium bromide (MTT) method with minor modifications (Feoktistova et al., 2016; Mosmann, 1983). Briefly, the cells were seeded in 96 well plates (100 μl/well at a density of 3x10^5^ cells/ml) and incubated for 24 h at 37 °C in 5%CO2. After 24 hours, cells were treated with various concentrations of the tested compounds in triplicates. 24 hours later, the supernatant was discarded, and cell monolayers were washed with sterile 1x phosphate buffer saline (PBS) 3 times and MTT solution (20 μl of 5 mg/ml stock solution) was added to each well and incubated at 37 °C for 4 hours followed by medium aspiration. In each well, the formed formazan crystals were dissolved with 200 μL of acidified isopropanol (0.04 M HCl in absolute isopropanol = 0.073 mL HCL in 50 mL isopropanol). The absorbance of formazan solutions was measured at λ max 540 nm with 620 nm as a reference wavelength using the Anthos Zenyth 200rt plate reader (Anthos Labtec Instruments, Heerhugowaard, Netherlands). The untreated control cells and blank optical density (OD) readings were used to normalize the OD readings from treated wells. The half-maximal cytotoxic concentration (CC_50_) was calculated using GraphPad Prism software using nonlinear regression analysis after plotting log concentrations versus normalized responses.

### Mode of Anti-SARS-CoV-2 Action

We aimed to investigate whether CPM, with a pre-determined anti-SARS-CoV-2 activity, can influence aspects of (a) viral adsorption, (b) viral replication, or (c) viricidal effect.

### Adsorption Mechanism

The viral adsorption mode of action was tested according to Zhang et al., with minor modifications (Zhang et al., 1995). Briefly, Vero E6 cells were cultivated in a six-well plate and incubated at 37 °C in a humidified 5 % CO_2_ incubator for 12-24 hours to reach 80-90% confluency. Subsequently, the growth media were aspirated, and the cell monolayers were washed with 1X PBS. Cell monolayers were treated with the indicated CPM concentrations (156, 78, 39, 19 μg/mL, in triplicate) at 4°C to minimize cellular drug uptake. One hour later, the applied drug inoculum was replaced with a countable dilution of the virus (Titer of stock virus = 7.5 x 10^5^ PFU/mL) for another 1 hour at 37 °C in a humidified 5 % CO_2_ incubator. After 1 hour of contact time, 3 mL of overlayer medium of DMEM supplemented with 2% agarose, 0.2 bovine serum albumin (BSA), and 1% antibiotic–antimycotic mixture was added to the cell monolayer. Uninfected control cells were included in each plate. Plates were incubated for three days and then were fixed using 10% formalin solution for 1 hour and stained with crystal violet. The percentages of viral inhibition were calculated based on untreated virus control wells.

### Replication Mechanism

The effect of each compound on the viral replication was studied as previously described (Kuo et al., 2002). Briefly, Vero E6 cells were cultivated in a six-well plate and incubated at 37 °C for 24 hours. The cells were infected with a countable dilution of the virus (titer of stock virus = 7.5 x 10^5^ PFU/mL) and then incubated for 1 hour at 37 °C for virus internalization or infection. The cells were washed using 1X PBS to remove the non-adsorbed virus. CPM (156, 78, 39, 19 μg/mL) was added to each well of infected cells. After 1 hour, cell monolayers were washed once with 1X PBS, and then 3 mL of overlayer medium of DMEM supplemented with 2% agarose and 1% antibiotic–antimycotic mixture was added to the cell monolayer. Uninfected control cells were included in each plate. Plates were incubated for three days and then were fixed using 10% formalin solution for 1 hour and stained with crystal violet. The percentages of viral inhibition were calculated based on untreated virus control wells.

### Virucidal Mechanism

The virucidal mechanism was assayed following a previously described protocol (Schuhmacher et al., 2003). In a 6-well plate, Vero E6 cells were cultivated (10^5^ cells/mL) for 24 h at 37 °C. An aliquot of 200 μL of serum-free DMEM containing stock SARS-CoV-2 (stock virus titer = 7.5 x 10^5^ PFU/mL) was added to the drug CPM at the predefined promising inhibitory concentrations. After 1 hour incubation, the mixture was 10-fold diluted three successive times using serum-free medium (7.5 x 10^2^ PFU/mL), which still allowed viral particles to grow on Vero-E6 cells at countable plaque forming units (PFU) with neglected residual drug concentration. Next, an aliquot of 100 μL of each dilution versus control untreated/diluted virus was added to the Vero E6 cell monolayers that are covered with 600 μL of serum-free DMEM. After 1 hour of contact time, a DMEM overlayer was added to the cell monolayer. Plates were left to solidify and incubated at 37 °C to allow the formation of viral plaques. The plaques were fixed and stained to calculate the percentage reduction in plaque formation. This value was compared to control wells comprising cells infected with the virus and not pretreated with the tested material.

### Chemoinformatics Molecular Modelling

The x-ray crystal structure coordinates of SAR-CoV-2 main protease (M^pro^) were retrieved from the Protein Data Bank (PDB) (PDB ID: 6yef and 6lu7) (Zhenming Jin et al., 2020) in addition to the retrieved receptor for S glycoprotein (PDB ID:6vsb) (Wrapp et al., 2020) with their co-crystallized bound ligand α-ketoamide, N3, and ligand 1 respectively. RNA polymerase of SARS-CoV-2 was retrieved from PDB (PDB ID:6m71). Furthermore, docking against ACE2 PDB (PDB ID:1r42) was modeled.

The docking study was performed using Open Eye scientific software version 2.2.5 (Santa Fe, NM (USA), http://www.eyesopen.com). For the validation of the docking study, the co-crystal bound ligands were docked. Both structures exhibited high similarity and overlaid each other as reported in our previous work (Allam et al., 2020).

### Physiochemical Parameter and Lipophilicity Calculations

Drug parameters including ClogP were calculated according to their practical values as reported in CHEMBL, Drug Bank, and PubChem free access websites. Lipinski’s rule (Rule of five) was calculated by the free access to the website https://www.molsoft.com/servers.html.

## Results

### Mechanism of Action

The half-maximal cytotoxic concentration (CC_50_) of the tested CPM in VERO E6 cells was 497.7 μg/ml (**Figure 1A**). The mode of anti-SARS-CoV-2 action indicated that CPM exerts dose-dependent antiviral effects via multiple mechanisms -direct-acting virucidal effect, viral replication inhibition, and viral adsorption inhibition (**Figure 1B**). The suggested model for the antiviral mode of CPM is shown in **Figure 6**.

**Figure 1.**
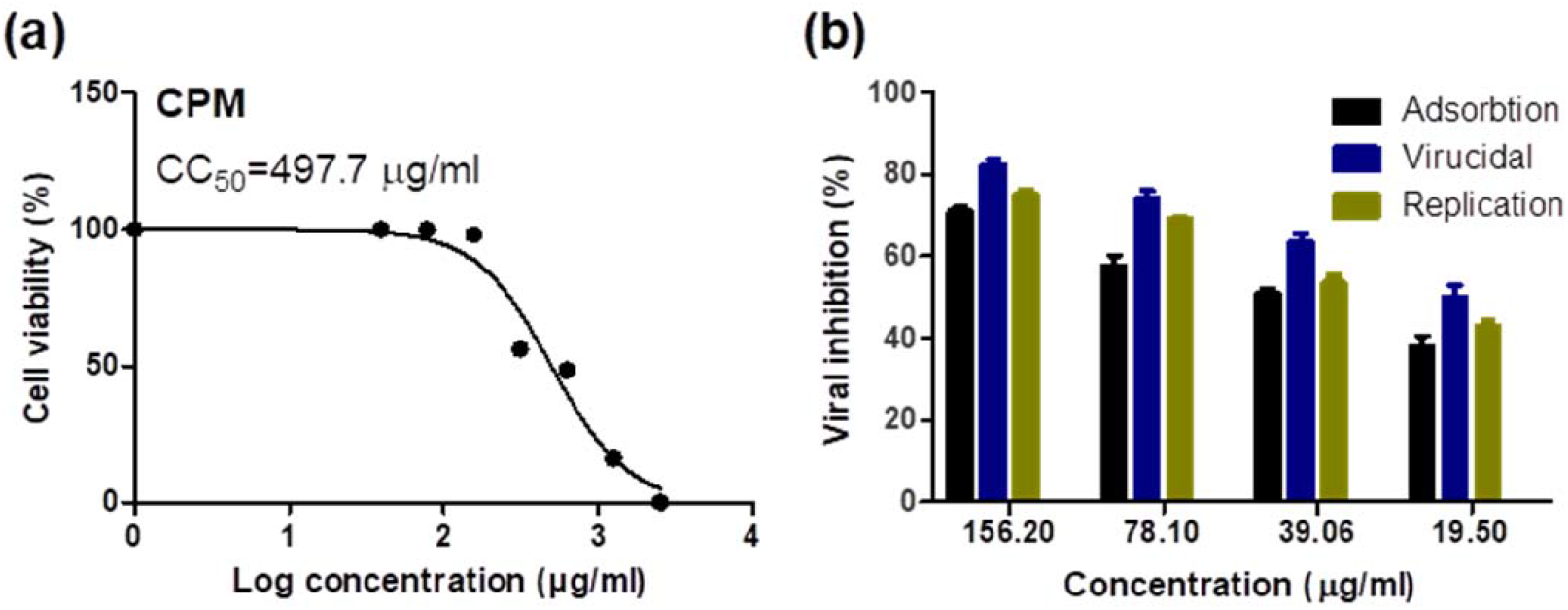
Cytotoxicity and cellular compartment mode of antiviral action of CPM in VERO E6 cells against SARS-CoV-2. (**A)** Cytotoxicity of the tested CPM in Vero E6 cells. The cytotoxicity of CPM based on the dose-response was determined using MTT. The 50% cytotoxic concentration (CC_50_) was calculated for each compound using nonlinear regression analysis of GraphPad Prism software (version 5.01). (**B)** Mode of anti-SARS-CoV-2 action of Chlorpheniramine Maleate. Virucidal, viral replication inhibition, and viral adsorption inhibition mechanisms were studied for CPM at different concentrations using plaque reduction assay.

### Molecular Docking Studies

The analysis of the chlorphenamine CPM’s binding mode formed interactions with the receptor of M^pro^ (PDB ID:6lu7) through hydrophobic-hydrophobic interaction (Figure 2a). From Figure 2b, CPM (green color) showed overlay with N3, the co-crystalized ligand, (grey color).

**Figure 2.**
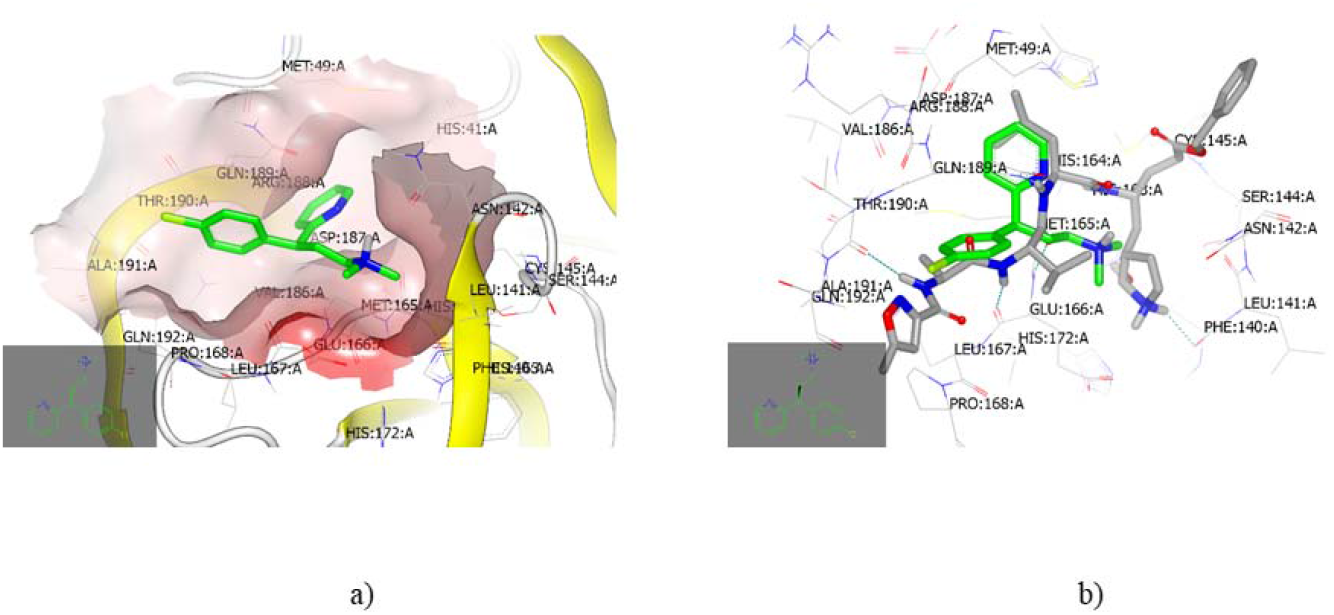
Molecular docking of Chlorpheniramine Maleate with SARS-CoV-2 Main Protease (M^pro^)A) Chlorpheniramine Maleate formed an interaction with the receptor of M^pro^ (PDB ID:6lu7) of SARS-CoV-2 through hydrophobic-hydrophobic interaction; B) CPM (green color) showed overlay with N3, the co-crystalized ligand M^pro^ (PDB ID:6lu7) of SARS-CoV-2, (grey color).

Docking of CPM with S glycoprotein (PDBID: 6vsb) presented hydrophobic-hydrophobic interaction (Figure 3a). The analysis of the binding mode and pose of this drug (green color) in comparison to the co-crystalized ligand (grey) is shown in Figure 3b. The *P*-chlorophenyl ring overlayed with *P*-methoxy moiety and the *N, N-dimethylethanolamine* part.

**Figure 3.**
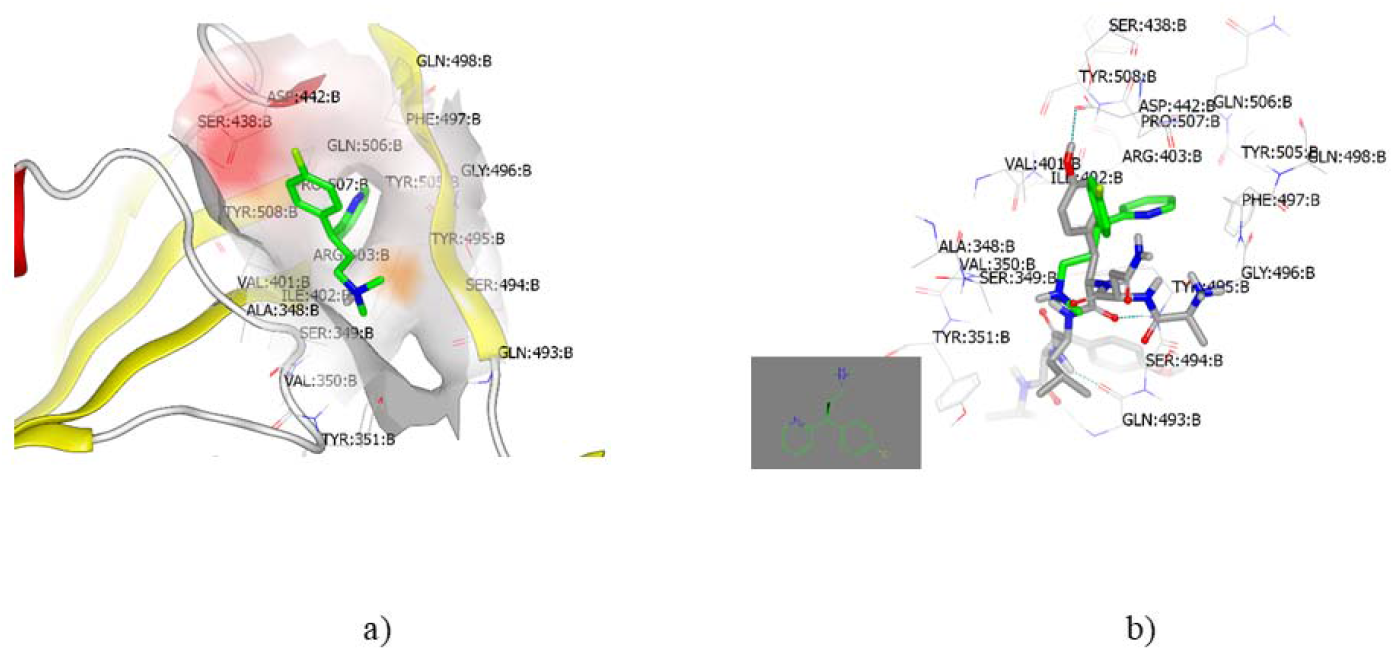
Molecular docking of Chlorpheniramine Maleate with SARS-CoV-2 spike protein. A) Chlorpheniramine Maleate formed an interaction with the receptor of spike protein (PDB ID:6vsb) of SARS-CoV-2 through hydrophobic-hydrophobic interaction; B) Chlorpheniramine Maleate (green color) showed overlay with the co-crystalized ligand spike protein (PDB ID:6vsb) of SARS-CoV-2, (grey color) with little similarity in binding pose and mode.

**Figure 4.**
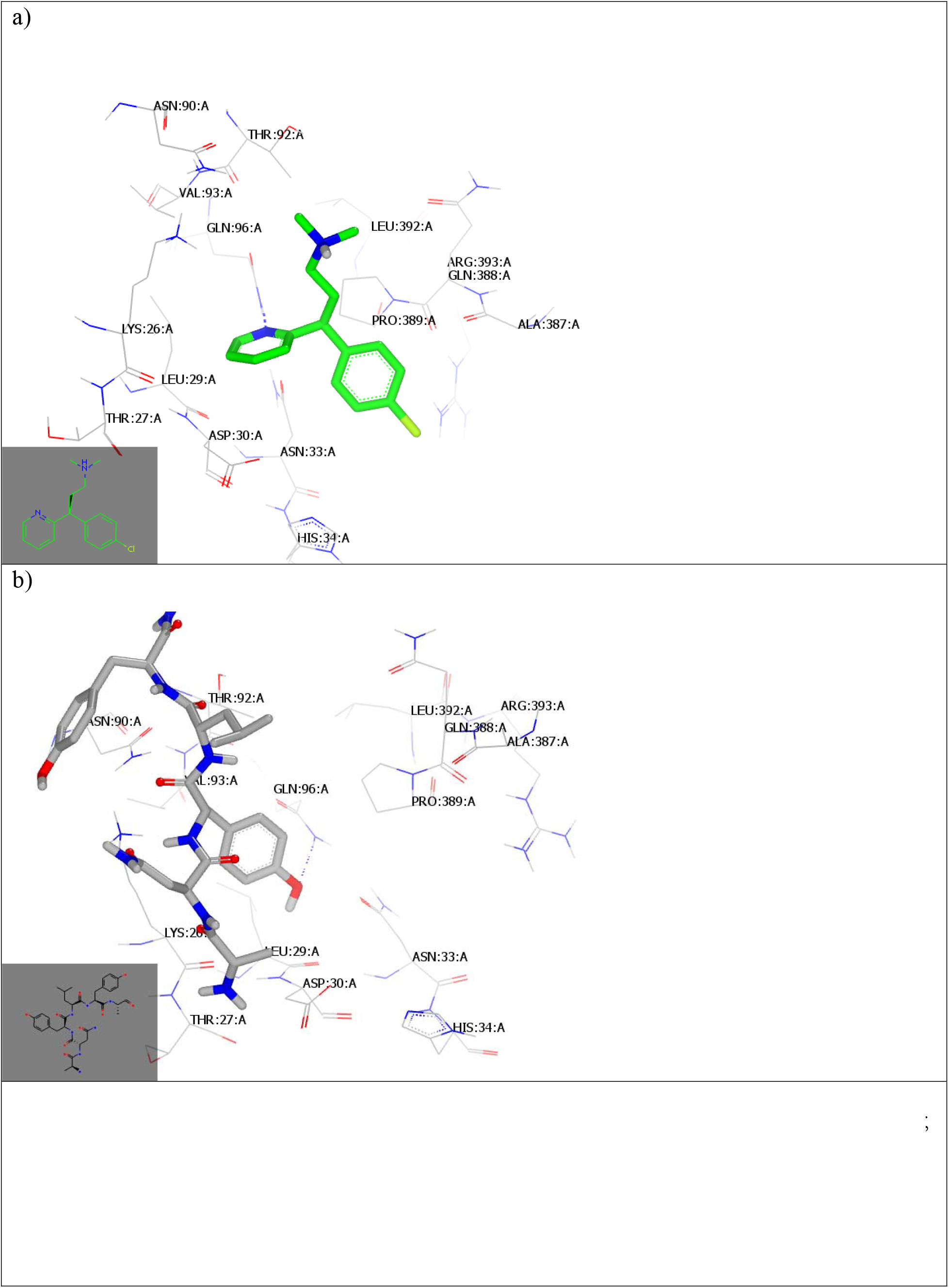
**a)** Molecular docking of Chlorpheniramine Maleate with ACE2 receptor (ID 1R42); b) Molecular docking of standard ligand for SARS-CoV-2 spike protein with ACE2 receptor (ID 1R42))

Docking with RNA polymerase (PDB ID:6m71 [4] showed that CPM formed a hydrogen bond (HB) with Asn:79 A through the nitrogen atom of the pyridine moiety. The ethylamine derivative is adopted deeply inside the pocket of the active site, Figure 5A.

**Figure 5.**
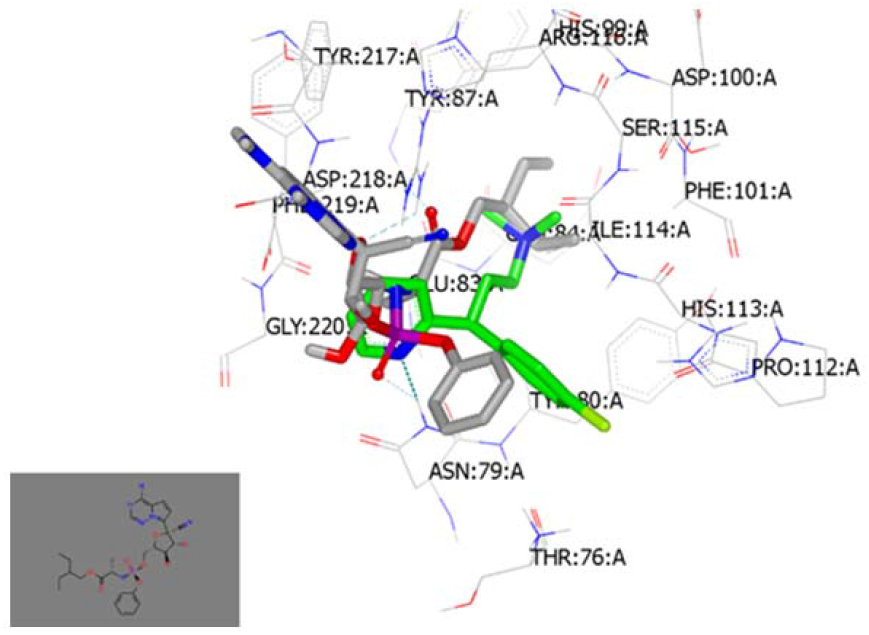
Molecular docking of Chlorpheniramine Maleate with SARS-CoV-2 RNA polymerase Visual representation using Vida application: a) Chlorpheniramine Maleate interacted with ID:6m71 active site by forming hydrogen bonds with Asn:79 A through the nitrogen atom of pyridine moiety.

**Figure 6.**
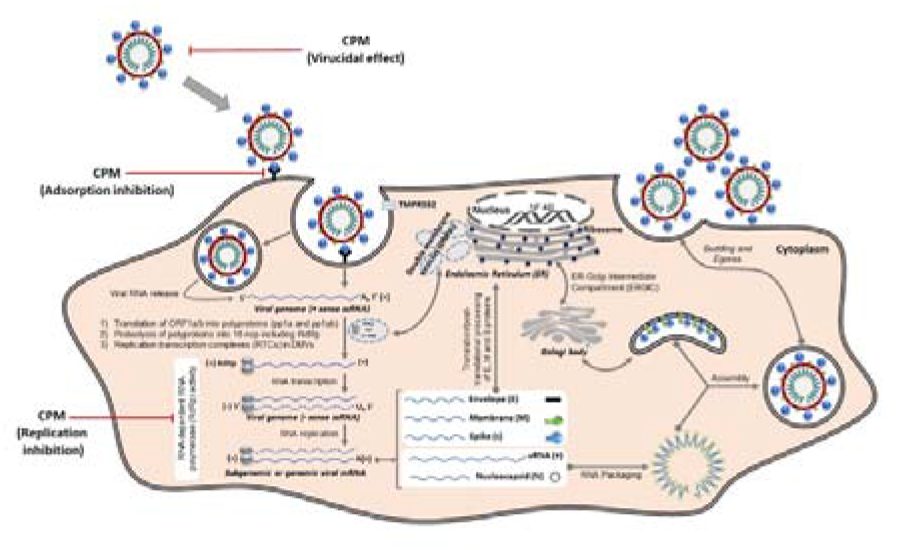
Model for SARS-CoV-2 antiviral mechanisms mediated by Chlorpheniramine Maleate adapted and modified from (Ahmed Mostafa et al., 2020).

ACE2 is a functional receptor that allows SARS-CoV-2 to enter human lung cells [via its spike (S) protein (Ahmad et al., 2021). Molecular docking with ACE receptors was then performed.

Docking of CPM with ACE2 receptor (PDB: ID1R42) showed that the NH of the pyridine ring participated in hydrogen bond (HB) formation with Gln 96 AA, figure 4a (Towler et al., 2004). However, CPM does not interact with the same domain as the co-crystallized ligand. The standard spike protein-ligand was also docked and illustrated HB formation with the same amino acid, figure 4b (Wrapp et al., 2020).

According to previous molecular docking and *in vitro* viral inactivation and molecular mechanisms, CPM can inhibit virus replication. In this regard, lipophilic metric studies will be implemented in comparison to the reported drug remdesivir (RNA polymerase inhibitor, antiviral replication). Lipophilic metric analysis, especially Ligand efficiency (LE) and Ligand lipophilic efficiency (LLE) scores, is very important in the drug-ability process and during the drug repositioning strategy.

The ligand and target interaction are the main parameters in the drug discovery and development pipeline. Presently, validation of the molecular size, and lipophilicity (CLogP) together with drug activities (pIC_50_), known. as “optimization measures” are required.

The challenge in drug discovery is to improve the activity while keeping lipophilicity constant to avoid any “molecular obesity” during drug development. An acceptable lead drug should have an LLE value ≥ 3 while an LLE value ≥ 5 is recommended for a drug-like candidate. As illustrated in Table 1, CPM displayed LE better than remdesivir (0.39, 0.17 respectively) while remdesivir illustrated LEE little value higher than CPM (2.64, 2.26) respectively. CPM can penetrate the blood-brain barrier (BBB) more than remdesivir as shown in Table 1. CPM has Lipinski rule of five lower than remdesivir.

**Table 1.**
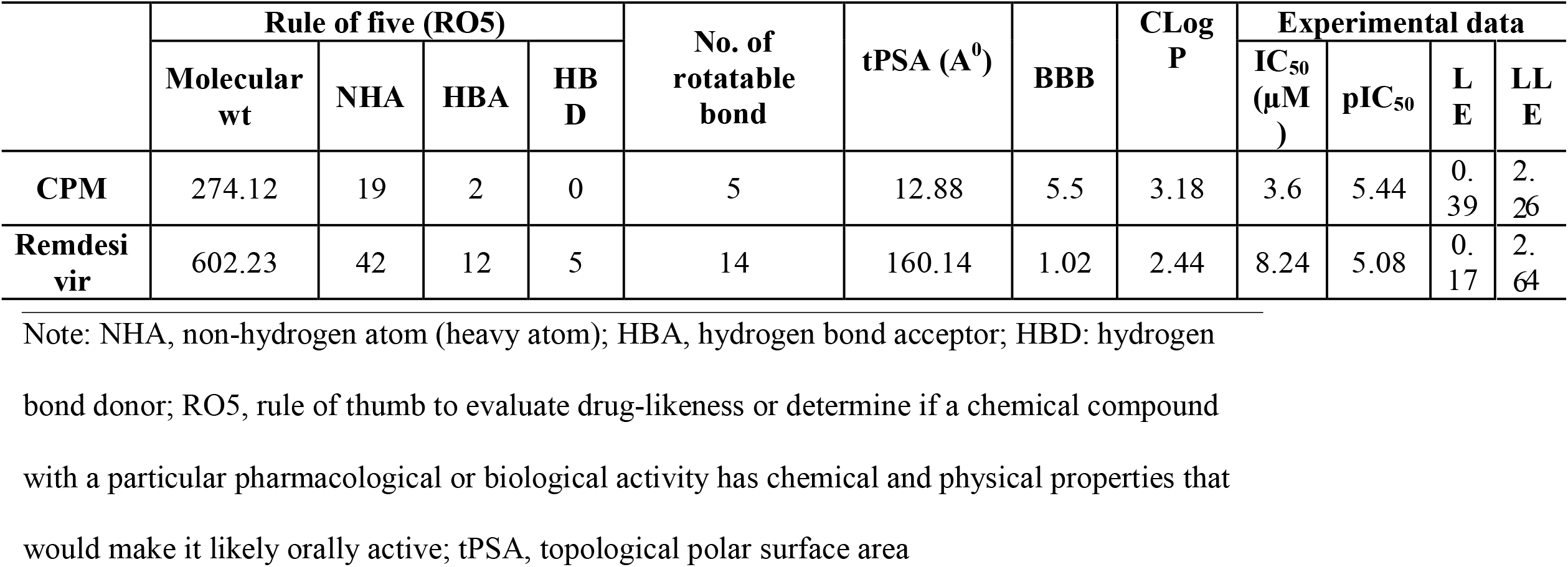
Selected physicochemical properties, and ligand efficiency scores of CP and remdesivir against SARS-CoV-2

## Discussion

Herein we investigated the impact of CPM treatment *in vitro* on the three main compartments during the SARS-CoV-2 replication cycle, including the direct-acting viricidal effect, antagonizing viral entry by interference with viral adsorption, and antagonizing viral replication inside the host cell. The main findings of the present study are that CPM can inhibit SARS-CoV-2 by affecting viral adsorption, replication, and direct virucidal effect. Chemoinformatic analysis revealed that CPM is a promising anti-SARS-CoV-2 with a multitarget effect against SARS-CoV-2. These data suggest that CPM has a potential broad-spectrum antiviral application. Mounting evidence suggests that CPM has both antiviral and anti-inflammatory actions that could be beneficial in treating COVID-19. Furthermore, molecular modeling data analyzed the antiviral activity of CPM, comparing the chemical structure of different over-the-counter drugs CPM was structurally like drugs known to have an anti-inflammatory and drugs known to have antiviral effect in drug-receptor interactions (Black, 2022; Gies et al., 2020). Clinical studies have shown that CPM improves clinical recovery and prevents hospitalizations in patients with COVID-19 (Black, 2022; Morán Blanco et al., 2021; Sanchez-Gonzalez et al., 2022). In a two-part clinical study conducted by our team, ACCROS-I, and ACCROS-II, suggested that CPM intranasal-administered is an effective early intervention to reduce symptoms and accelerate clinical recovery in COVID-19 patients (Valerio-Pascua et al., 2022). Results showed that there were less hospitalization and severe cases. The study provides evidence of the main mechanisms accountable for the antiviral actions of CPM against SARS-CoV-2 and based on the results, it is anticipated that CPM could significantly decrease viral load in patients affected by COVID-19.

Currently, in the market, two major medications received emergency use authorization (EUA) from the FDA for the treatment of COVID-19 Paxlovid and remdesivir. Paxlovid is an oral antiviral medication intended for patients with mild to moderate COVID-19, and Remdesivir is an intravenous antiviral for the treatment of both hospitalized and non-hospitalized COVID-19 patients. In December 2022, Pfizer announced that Paxlovid did not look likely to meet its main goal of alleviating COVID-19 Symptoms. This announcement raised concern about the potential increase in hospitalizations and the severity of cases. With prior clinical evidence and in vitro studies, it is plausible that CPM administered intranasally can decrease clinical symptomatology and accelerate clinical recovery. In reducing the hospitalization rate and severity of COVID-19 cases, we believe that early intervention aiming at the reduction of symptoms and time to recovery is a significant unmet need for treatment options to help combat this disease.

Furthermore, CPM has been shown to display broad-spectrum antiviral effects, including influenza, Ebola, and various strains of SARS-CoV-2 (Ekins et al., 2014; Xu et al., 2018; Westover et al., 2020; Sanchez-Gonzalez et al., 2022). However, the present study focused on viral targets and not host-directed targets. The authors cannot discard the possibility that other mechanisms, including host-targeted effects, may be in play as part of CPM’s antiviral effects needing further transcriptomic and proteomic studies.

## Conclusion

CPM can inhibit SARS-CoV-2 by affecting viral adsorption, replication, and direct virucidal effect. These data suggest that CPM has a potential broad-spectrum antiviral application. Furthermore, this study highlights the necessity for launching clinical studies for using CPM in combination or prior administration with other drugs in COVID-19 patients. Whether CPM may exert its action via other host-targeted mechanisms warrants further investigation.

## Credit authorship contribution statement

All the authors, Yaseen A. M. M. Elshaier, Ahmed Mostafa, Fernando Valerio-Pascua, Mari L. Tesch, Joshua M. Costin, Franck F. Rahaghi have contributed to the Methodology, Validation, Formal analysis, Writing -original draft, Writing -review & editing.

## References

Ahmad, I., Pawara, R., Surana, S., and Patel, H. (2021). The Repurposed ACE2 Inhibitors: SARS-CoV-2 Entry Blockers of Covid-19. Top Curr Chem (Cham) 379, 40.

Allam, A. E., Assaf, H. K., Hassan, H. A., Shimizu, K., & Elshaier, Y. A. M. M. (2020). An in silico perception for newly isolated flavonoids from peach fruit as privileged avenue for a countermeasure outbreak of COVID-19 [10.1039/D0RA05265E]. RSC Advances, 10(50), 29983–29998. 10.1039/D0RA05265E

Black, S. (2022). Molecular Modeling and Preliminary Clinical Data Suggesting Antiviral Activity for Chlorpheniramine (Chlorphenamine) Against COVID-19. Cureus, 14(1), e20980. 10.7759/cureus.20980

Feoktistova, M., Geserick, P., & Leverkus, M. (2016). Crystal Violet Assay for Determining Viability of Cultured Cells. Cold Spring Harb Protoc, 2016(4), pdb.prot087379. 10.1101/pdb.prot087379

Gies, V., Bekaddour, N., Dieudonné, Y., Guffroy, A., Frenger, Q., Gros, F., Rodero, M. P., Herbeuval, J. P., & Korganow, A. S. (2020). Beyond Anti-viral Effects of Chloroquine/Hydroxychloroquine. Front Immunol, 11, 1409. 10.3389/fimmu.2020.01409

Jabeen, I., Pleban, K., Rinner, U., Chiba, P., & Ecker, G. F. (2012). Structure-activity relationships, ligand efficiency, and lipophilic efficiency profiles of benzophenone-type inhibitors of the multidrug transporter P-glycoprotein. J Med Chem, 55(7), 3261–3273. 10.1021/jm201705f

Jin, Z., Du, X., Xu, Y., Deng, Y., Liu, M., Zhao, Y., Zhang, B., Li, X., Zhang, L., Peng, C., Duan, Y., Yu, J., Wang, L., Yang, K., Liu, F., Jiang, R., Yang, X., You, T., Liu, X., … Yang, H. (2020). Structure of M(pro) from SARS-CoV-2 and discovery of its inhibitors. Nature, 582(7811), 289–293. 10.1038/s41586-020-2223-y

Jin, Z., Du, X., Xu, Y., Deng, Y., Liu, M., Zhao, Y., Zhang, B., Li, X., Zhang, L., Peng, C., Duan, Y., Yu, J., Wang, L., Yang, K., Liu, F., Jiang, R., Yang, X., You, T., Liu, X., … Yang, H. (2020). Structure of Mpro from SARS-CoV-2 and discovery of its inhibitors. Nature, 582(7811), 289–293. 10.1038/s41586-020-2223-y

Kandeil, A., Mostafa, A., El-Shesheny, R., Shehata, M., Roshdy, W.H., Ahmed, S.S., Gomaa, M., Taweel, A.E., Kayed, A.E., Mahmoud, S.H., Moatasim, Y., Kutkat, O., Kamel, M.N., Mahrous, N., Sayes, M.E., Guindy, N.M.E., Naguib, A., and Ali, M.A. (2020). Coding-Complete Genome Sequences of Two SARS-CoV-2 Isolates from Egypt. Microbiol Resour Announc 9.

Kuo, Y. C., Lin, L. C., Tsai, W. J., Chou, C. J., Kung, S. H., & Ho, Y. H. (2002). Samarangenin B from Limonium sinense suppresses herpes simplex virus type 1 replication in Vero cells by regulation of viral macromolecular synthesis. Antimicrob Agents Chemother, 46(9), 2854–2864. 10.1128/aac.46.9.2854-2864.2002

Lee, R. J., Kofonow, J. M., Rosen, P. L., Siebert, A. P., Chen, B., Doghramji, L., Xiong, G., Adappa, N. D., Palmer, J. N., Kennedy, D. W., Kreindler, J. L., Margolskee, R. F., & Cohen, N. A. (2014). Bitter and sweet taste receptors regulate human upper respiratory innate immunity. J Clin Invest, 124(3), 1393–1405. 10.1172/jci72094

Ligand efficiency as a guide in fragment hit selection and optimization. (2010). Drug Discov Today Technol, 7(3), e147–202. 10.1016/j.ddtec.2010.11.003

Morán Blanco, J. I., Alvarenga Bonilla, J. A., Homma, S., Suzuki, K., Fremont-Smith, P., & Villar Gómez de Las Heras, K. (2021). Antihistamines and azithromycin as a treatment for COVID-19 on primary health care -A retrospective observational study in elderly patients. Pulm Pharmacol Ther, 67, 101989. 10.1016/j.pupt.2021.101989

Mosmann, T. (1983). Rapid colorimetric assay for cellular growth and survival: application to proliferation and cytotoxicity assays. J Immunol Methods, 65(1-2), 55–63. 10.1016/0022-1759(83)90303-4

Mostafa, A., Kandeil, A., Shehata, M., El Shesheny, R., Samy, A. M., Kayali, G., & Ali, M. A. (2020). Middle East Respiratory Syndrome Coronavirus (MERS-CoV): State of the Science. Microorganisms, 8(7), 991. https://www.mdpi.com/2076-2607/8/7/991

Mostafa, A., Kandeil, A., Y, A. M. M. E., Kutkat, O., Moatasim, Y., Rashad, A. A., Shehata, M., Gomaa, M. R., Mahrous, N., Mahmoud, S. H., GabAllah, M., Abbas, H., Taweel, A. E., Kayed, A. E., Kamel, M. N., Sayes, M. E., Mahmoud, D. B., El-Shesheny, R., Kayali, G., & Ali, M. A. (2020). FDA-Approved Drugs with Potent In Vitro Antiviral Activity against Severe Acute Respiratory Syndrome Coronavirus 2. Pharmaceuticals (Basel), 13(12). 10.3390/ph13120443

Reznikov, L. R., Norris, M. H., Vashisht, R., Bluhm, A. P., Li, D., Liao, Y. J., Brown, A., Butte, A. J., & Ostrov, D. A. (2021). Identification of antiviral antihistamines for COVID-19 repurposing. Biochem Biophys Res Commun, 538, 173–179. 10.1016/j.bbrc.2020.11.095

Rizvi, A. A. S., Ferrer, G., Khawaja, A. U., & Sanchez-Gonzalez, A. M. (2022). Chlorpheniramine, an Old Drug with New Potential Clinical Applications: A Comprehensive Review of the Literature. Current Reviews in Clinical and Experimental Pharmacology, 17, 1–1. 10.2174/2772432817666220601162006

Sanchez-Gonzalez, M., Rizvi, S. A., Torres, J., & Ferrer, G. (2021). A Randomized Controlled Pilot Trial to Test the Efficacy of Intranasal Chlorpheniramine Maleate With Xylitol for the Treatment of Allergic Rhinitis. Cureus, 13(3), e14206. 10.7759/cureus.14206

Sanchez-Gonzalez, M. A., Westover, J. B., Rizvi, S. A. A., Torres, J., & Ferrer, G. A. (2022). Intranasal Chlorpheniramine Maleate for the treatment of COVID-19: Translational and Clinical Evidence. Medical Research Archives, 10(3). 10.18103/mra.v10i3.2752

Schuhmacher, A., Reichling, J., & Schnitzler, P. (2003). Virucidal effect of peppermint oil on the enveloped viruses herpes simplex virus type 1 and type 2 in vitro. Phytomedicine, 10(6-7), 504–510. 10.1078/094471103322331467

Torres, J., Go, C., Camacho, G., Sanchez-Gonzalez, M., & Ferrer, G. (2021). Chlorpheniramine Maleate Nasal Spray In COVID-19 Patients: Case Series. J Clin Exp Pharmacol, 10(2), 3. 10.35248/2161-1459.21.10.275

Towler, P., Staker, B., Prasad, S.G., Menon, S., Tang, J., Parsons, T., Ryan, D., Fisher, M., Williams, D., Dales, N.A., Patane, M.A., and Pantoliano, M.W. (2004). ACE2 X-ray structures reveal a large hinge-bending motion important for inhibitor binding and catalysis. J Biol Chem 279, 17996–18007.

Valerio-Pascua, F., Mejia, E.J.P., Tesch, M., Godoy, J., Fuentes, C.L., Erazo, G., Bermúdez, M., Pineda, M.F.V., Rivzi, S.a.A., Cabrera, A., Chauhan, Z., Grullón-Franco, S., Paulino-Then, J., Garcia, N., Williams, J., and Rahaghi, F. (2022). “Chlorpheniramine Intranasal Spray to Accelerate COVID-19 Clinical Recovery in an Outpatient Setting: The ACCROS Trials”. Research Square).

Westover, J. B., Ferrer, G., Vazquez, H., Bethencourt-Mirabal, A., & Go, C. C. (2020). In Vitro Virucidal Effect of Intranasally Delivered Chlorpheniramine Maleate Compound Against Severe Acute Respiratory Syndrome Coronavirus 2. Cureus, 12(9), e10501. 10.7759/cureus.10501

Wrapp, D., Wang, N., Corbett, K. S., Goldsmith, J. A., Hsieh, C.-L., Abiona, O., Graham, B. S., & McLellan, J. S. (2020). Cryo-EM structure of the 2019-nCoV spike in the prefusion conformation. Science,367(6483), 1260–1263. doi:10.1126/science.abb2507

Xu, W., Xia, S., Pu, J., Wang, Q., Li, P., Lu, L., & Jiang, S. (2018). The Antihistamine Drugs Carbinoxamine Maleate and Chlorpheniramine Maleate Exhibit Potent Antiviral Activity Against a Broad Spectrum of Influenza Viruses. Front Microbiol, 9, 2643. 10.3389/fmicb.2018.02643

Zhang, J., Zhan, B., Yao, X., Gao, Y., & Shong, J. (1995). [Antiviral activity of tannin from the pericarp of Punica granatum L. against genital Herpes virus in vitro]. Zhongguo Zhong Yao Za Zhi, 20(9), 556–558, 576, inside backcover.

